# A Structured Professional Development Curriculum for Postdoctoral Fellows Leads to Recognized Knowledge Growth

**DOI:** 10.1101/2020.10.15.340059

**Authors:** Kaylee Steen, Jay Vornhagen, Zara Y. Weinberg, Julie Boulanger-Bertolus, Arvind Rao, Margery Evans Gardner, Shoba Subramanian

## Abstract

Postdoctoral training enables research independence and professional readiness. National reports have emphasized professional development as a critical component of this training period. In response, many institutions are establishing transferable skills training workshops for postdocs; however, the lack of structured programs and an absence of methods to assess outcomes beyond participant satisfaction surveys are critical gaps in postdoctoral training. To address these shortcomings, we took the approach of structured programming and developed a method for controlled assessment of outcomes. Our program You^3^ (You, Your Team, Your Project), co-designed by postdoctoral fellows, focused on a structured array of management and leadership skills agnostic of ultimate career path(s). We then measured outcomes in a controlled manner, by systematically comparing perceived knowledge and growth of participants with non-participants as the control group. You^3^ participants discern greater growth, independent of number of years in training, in competencies overall compared to the control group. This growth was shown by multiple criteria including self-reporting and associative analysis. Correspondingly, You^3^ participants reported greater knowledge in 75% of the modules when compared to controls. These data indicate that structured learning, where postdocs commit to a curriculum via a cohort-structure, leads to positive outcomes and provides a framework for programs to assess outcomes in a rigorous manner.

## Introduction

Postdoctoral training is a unique period that is defined by the National Postdoctoral Association (nationalpostdoc.org) as “temporary time for scholarly work as well as acquisition of professional skills for career success”. To be successful in their next career step, postdoctoral fellows (referred to as ‘postdocs’ for the remainder of this article) need to develop a wide variety of skills, including technical and non-technical skills. Although the importance of technical skills has been recognized for many years, growing evidence shows non-technical skills, also known as transferable skills, are also vital to succeeding as a professional during and beyond training (1–5). A well-rounded portfolio is especially important in current times, where less than 25% of the PhD graduates move on to a tenure track (TT) position, while the rest pursue a distinct array of individual paths (6).

The unique nature of the postdoctoral training period presents several challenges for developing transferable skills. Postdocs are hired by individual faculty advisors and do not typically enter the academic system as part of a cohort. They start their appointment asynchronously, and the orientation, onboarding routines, and access to non-research resources are highly variable within and across academic units. Postdoctoral training is largely apprentice-based, leading to disparities in the scope and breadth of training, depending on their advisor’s experience and encouragement in combination with institutional support for expanded learning opportunities. The lack of structured professional skill development opportunities underserve the nearly 80% of PhD students that go on to do postdoctoral training (7). Many institutions face these limitations (8), due to a combination of constraints on incentives, resources, personnel, and budget. Even in the case where there are more professional development resources to build programs, both access to and utilization of these programs vary heavily depending on the individual trainee’s circumstances.

Two key factors that are critical limitations in training postdocs are a lack of structured programs for standardized training and a lack of methods to assess outcomes scientifically. Research institutions are investing resources at the university, college, and departmental levels to provide seminars and workshops often based on the postdoc core competency guidelines set by the National Postdoctoral Association. Although well intentioned, these programs often follow an *ad hoc* pattern of topics and schedules with little structure, accountability, or evaluation of efficacy. A lack of a sequential structure leads to discontinuous learning, which can be ineffective (9). Moreover, the absence of a cohort or peer structure for postdocs compounds shortcomings within these programs, leading to loss of peer-community groups and peer learning with a higher impact on historically excluded groups (10,11). Often, professional development programs are segregated by academic or non-academic career paths. This segregation makes it difficult for those who are undecided to choose the right program for them. These career-goal based professional development programs also lose the benefit of exchange and discourse between individuals of different mindsets and backgrounds. The lack of structure is beginning to be addressed by systems like the NIH BEST program (12) which recognizes the vital role of a robust infrastructure for professional development in scientific and science-related careers. However, even when institutes put in place structured professional development programs, outcomes are primarily assessed using basic methods such as satisfaction surveys. A lack of rigorous assessments for professional development program efficacy is pervasive across all types of skill building programs partly because of no useful mechanism(s) for conventional formative or summative assessments. This gap is also driven by the non-technical nature of the topics covered, which are challenging to quantify in advanced learners. As a result, we have no clear and controlled assessment methods that measure impact via participants’ knowledge and growth in the topic in a systematic manner.

To effectively address these key limitations, we developed and implemented an innovative leadership and management program titled “You^3”^. Our structured program, co-designed and co-created by four postdoctoral fellows, used a cohort-based and sequential curriculum followed by measuring participants’ perceived knowledge and growth in a controlled method. Our results show that participants increased their own perceptions of current knowledge and growth in several key transferable skills that were measured against a control population who did not participate in our program. We propose You^3^ as a scalable framework that can seed and establish similar opportunities for postdocs in other institutions across the nation.

## Materials and Methods

### Needs Assessment Data Collection

#### Employer survey

The Office of Graduate and Postdoctoral Studies had commissioned miLEAD, a trainee-run non-profit consulting group on campus, to better understand career preparation and job market trends for graduate and postdoc learners. Based on data collected and analyzed by miLEAD, we compiled a list of the top skills hiring managers look for in applicants along with the perceived strengths and weaknesses Ph.D. holding candidates typically display in relation to these skills. miLEAD acquired this data by conducting interviews with multiple employment sectors including Academia, Biotech-Pharma, Legal-IP, Tech Transfer, Consulting, Writing, Government, and Bioinformatics. The data were further supported by analysis of 192 job postings to identify the list of required and preferred applicant skills, experience and attributes. miLEAD also identified the training needs of postdocs through a series of focus groups (n=18) and surveys (n=237) sent to University of Michigan Medical School (UMMS) postdocs. The postdoc needs data are represented via a summary of miLEAD group’s key findings.

#### UMMS Faculty survey

Before program planning, we sent an anonymous needs assessment survey to training faculty at the University of Michigan Medical School. The survey asked faculty to select all skills that applied to the question “Which transferable skills do you wish you had developed further prior to becoming a faculty member?”. Of the 266 respondents, 32.7% were Professors, 26.6% were Assistant Professors, 21.4% were Associate Professors, 5.2% were Clinical Instructional Faculty, 3% were Lecturers, 1.5% were Instructors, and 9.3% were Other. Faculty were able to select more than one skill in their response, and we counted the total number of times each topic was selected across each faculty category.

### You^3^ participant enrollment

Advertisement material of the You^3^ program was widely distributed to UMMS postdoctoral fellows via emails and flyers. Prospective applicants were asked to reflect on their professional experience and goals in order to assess their intentions and commitment to the You^3^ program. The questions were intentionally broad as to avoid self-selecting for postdoctoral fellows that were interested in a specific career trajectory over another. Applicants were also asked their career interest and were able to select as many options as were applicable. Supplemental Figure S1 displays complete application requirements. For reviewing the applications, we utilized a scale of 1-5 with 5 being excellent alignment between the applicant’s goals and the learning goals of You^3^ and 1 being low alignment. Applicants with an average score of 4 or higher were automatically accepted; applicants with a score between 3 and 4 were discussed by the committee, and a select few were requested to interview to better gauge their overall commitment and interest in the program. By this process, 33 of 35 applications were sent an acceptance letter into the program. 32 accepted the offer and committed to participation.

### Legend: Supplemental Figure 1: Application Questionnaire

*This application via Google forms was sent to all UMMS Postdocs to solicit participants*.

### Self-assessment pre-course survey to You^3^ participants

Prior to starting the You^3^ program, participants were surveyed on their training history in the core competencies of the program. Their responses were categorized into formal training (led by an instructor with engagement activities, i.e. workshop, class), informal training (Purely self-driven, i.e. online webinar, reading, etc.), and no training prior to enrollment.

### Self-assessment outcomes survey to control and participant groups

As a control group, all UMMS postdoctoral fellows received an online Qualtrics survey (IRB Exempt #HUM00172637) to anonymously assess their self-reported knowledge and growth in the eight topics covered by the You^3^ program (Fig S2). Based on each participant’s response, the survey items were binned according to the trainee’s prior experience with any UM skills building workshop, no skills building workshop experience, or participation in the You^3^ program. The following distribution of survey participants completed the questionnaire: You^3^ participants (n=23); control group who participated in some kind of professional development programming at UM (n=18); and did not participate in relevant programming (n=35).

### Legend: Supplemental Figure 2: Survey questions

*Survey questions that were sent out to You3 participants to measure their self-reported knowledge and growth around the eight modules after completing the program. This was compared to the control survey sent out to UMMS postdoctoral fellows that did not participate in the You*^*3*^ *program and were asked to report their knowledge and growth on the same metrics and time frame of the You*^*3*^ *program*.

### Statistical analysis for self-reported knowledge and growth metrics

For knowledge metric analysis, confidence score values were compared using Kruskal-Wallis test followed by Dunn’s posthoc test, and Pearson correlation coefficients and P values were calculated for each category and metric. For growth metric analysis, text-based respondent data was converted into numerical data, wherein 4 categories (no growth, low growth, medium growth, high growth) were converted to numerical values (1-4, respectively). Odds ratios and associated 95% confidence intervals, as well as Pearson correlation coefficients and P values, were calculated for each category and metric. For reliability analysis, Cronbach’s alpha was calculated for each question block (Growth and Current Knowledge) in R version 4.0.2 using the “psych” package. Kruskal-Wallis test followed by Dunn’s posthoc test were performed using GraphPad Prism version 8, and Pearson correlation coefficient and P-value calculations were performed and visualized in R version 4.0.2 using the “Hmisc” package.

## Results

To establish a comprehensive program, we focused on four achievable goals: (a) addressing specific needs of postdocs immediately transitioning into the workforce, (b) using a holistic view of professional development over eight consecutive weeks, (c) developing a cohort model toward fostering community, and (d) enabling management and leadership skills agnostic of ultimate career choice. At the end of the program, we developed an instrument to measure the impact of our program in a controlled manner by comparing participants’ against control non-participants in self-efficacy via knowledge and perceived value calibrated by growth in topic areas.

First, we used evidence-based data to determine the core topic list. We factored in needs assessments and workforce needs from the perspective of three key stakeholder groups: a) employers hiring PhD holders, b) UM Medical School (UMMS) faculty, and c) postdocs at our school during the time of survey.

We focused on the needs of employers by identifying deficiencies in professional skills for PhD job applicants based on interviews with 36 industry professionals and analysis of 192 non-academic job postings (data collected by miLEAD Consulting Group, Fig 1A). From the top skills that emerged, communication, teamwork and management (project, time, personnel, and budgeting) were perceived as weaknesses of PhD degree holders seeking employment (Fig 1A). These data were complemented by an anonymous survey distributed to UMMS faculty. We asked current training faculty in our college to identify one or more of 15 transferable skills that they were not proficient in prior to becoming a faculty member. Data from the 266 responses pointed at many common skills that were identified as deficiencies prior to assuming their faculty roles. Starting with the most frequently indicated, these skills included Grant/Scientific Writing, Team Building/Mentorship, Conflict Resolution, Statistics, Lab Business Management, and Lab Finances/Budgeting (Fig 1B). These data include responses from Assistant, Associate and Full Professors, and the results are consistent between all tracks. Finally, we compared the skills needed for successful careers to the gaps in training identified by current postdocs. Based on interviews and surveys collected from miLEAD Consulting, the current state of training was not deemed sufficient in preparing postdocs for professional careers, and the desired state was summarized in Figure 1C. These collective data from trainees, employers, and faculty are consistent with published work on PhD-holding employees (5)

**Figure 1:**
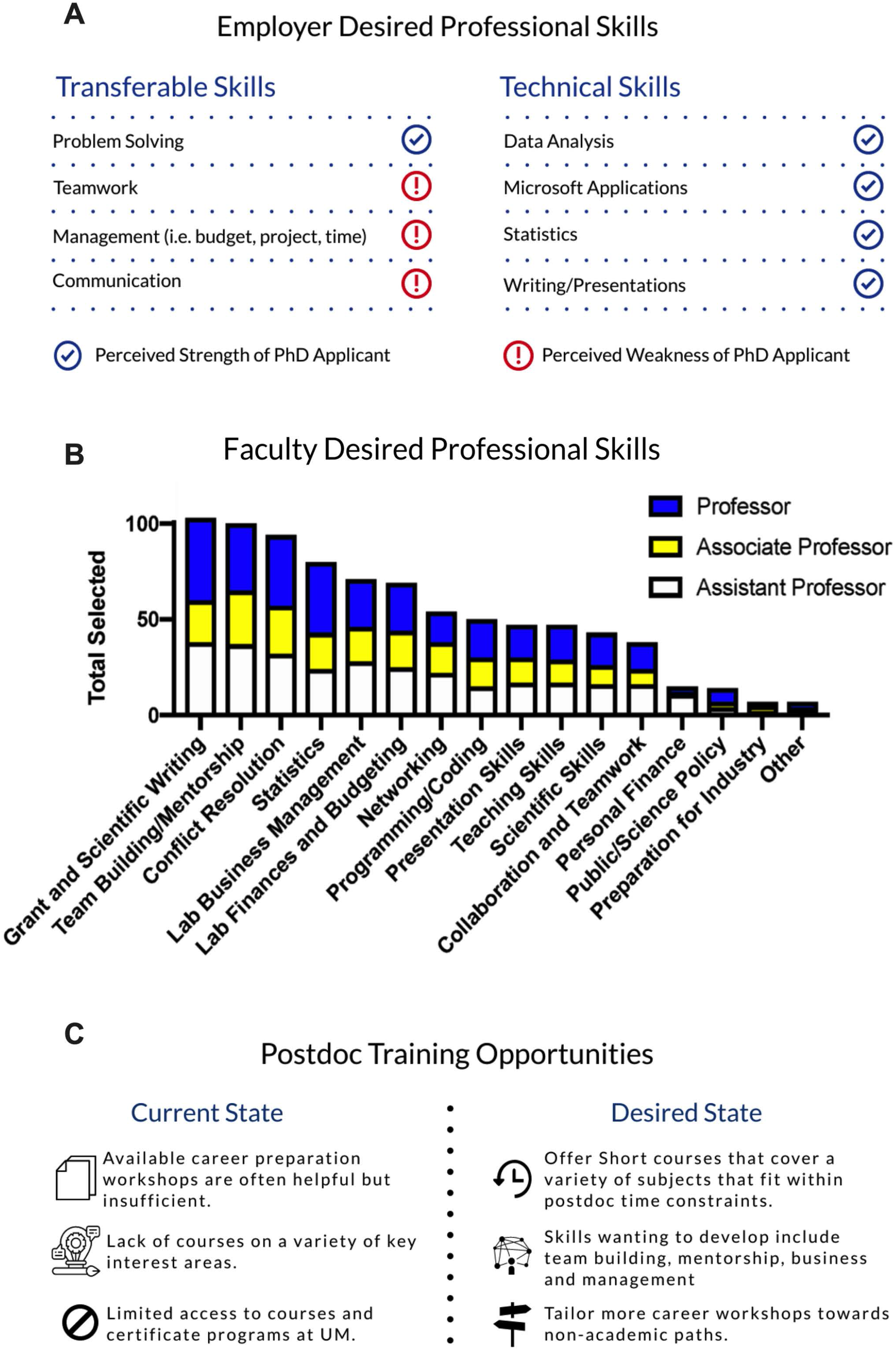
Postdocs do not receive adequate professional training based on needs assessment surveys and interviews. (A) miLEAD Consulting Group interviewed 36 industry professionals and analyzed 192 non-academic job postings to identify the professional skills employers seek in applicants. Those desired skills were separated into transferable and technical skills and marked as perceived weaknesses and strengths of PhD applicants based on industry interviews. (B) A needs assessment survey was distributed to UMMS faculty (266 responded), who rated the value each skill contributed to their career success. The faculty surveyed represented tenure-track (assistant, associate, and full professor), non-tenure track (lecturer and instructors), clinical-instructional track, and “other” category. For clarity, shown are responses from full, associate, and assistant professors, but the results were consistent across all faculty categories. (C) miLEAD Consulting Group served and interviewed current and former postdocs from 14 departments within UMMS in order to determine the professional development training postdocs receive. Their responses were analyzed and summarized into three major areas for their current and desired state of professional development training.

The needs assessment above fueled many robust discussions and literature searches among the steering committee, which consisted of four postdoc co-designers, leading to a robust program methodology as seen in Fig 2A. These data and discussions led us to focus on eight major topics that met the criteria of priorities for postdocs while being career agnostic. These topics were organized into three themes: Managing Yourself, Managing Your Project, and Managing Your Team (Figure 2B). This process also resulted in branding the program as *You*^*3*^: *Leadership and Management Program for Postdocs*: <u>You</u>, <u>You</u>r Team, <u>You</u>r Project (Fig 2B).

**Figure 2:**
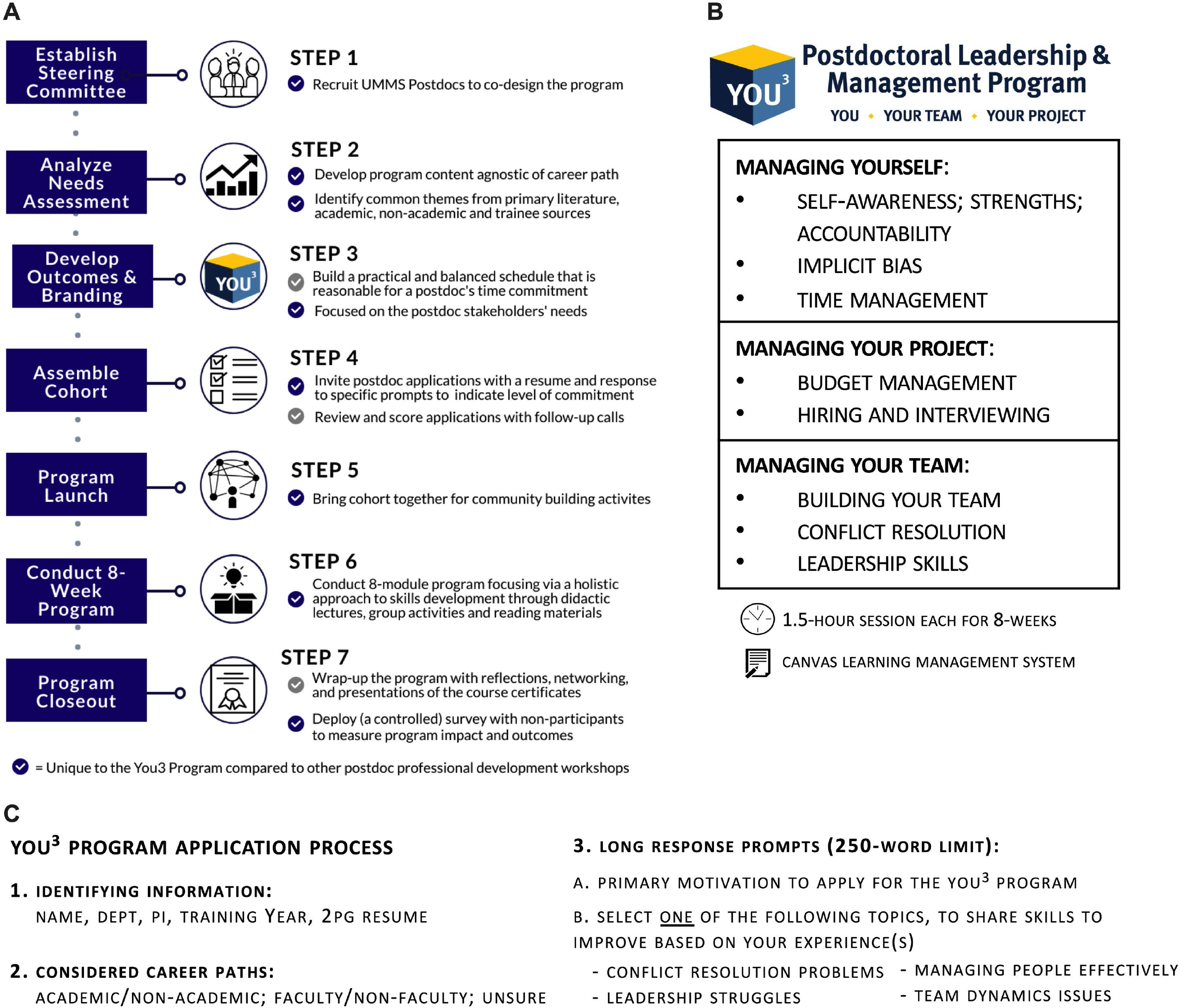
Overview of You^3^ program planning, framework, and application process. (A) Overview of the nine steps of planning, advertising, execution, and outcomes measurement of the You^3^ program. (B) Based on combined needs assessment and survey data, the You^3^ program content addressed 3 major themes: Managing Yourself, Managing Your Project, and Managing Your Team. Topics were then organized by theme, and each one was presented over the 8-week program in the order indicated in the table. (C) Responders to the prompts shown were then randomly divided into two groups, and each applicant group was evaluated by three members of the You^3^ steering committee.

We formally launched the inaugural You^3^ program in September 2019 using email marketing to postdoctoral fellows in UMMS and posted flyers across our research buildings. An open invitation intentionally queried diverse motivations for engaging in this program as evident in the application prompts and selection protocol (Fig 2C and Fig S1). Applicants were required to submit a two-page resume and respond to one of four simple prompts on the topics of conflict resolution, interpersonal and team-work experiences, leadership, and project management (Fig 2C). The application material was used during the selection process to their gauge interest and commitment. We also gathered data on applicants’ current career plans and prior training in the program’s core competencies. As seen in Figure 3A, out of 35 applicants, 19 were certain about one career path, whereas the remaining 21 chose more than one potential path. A pre-program anonymous survey indicated that the majority of applicants also reported they had little to no formal training in the skills covered in the You^3^ program, further highlighting the gap in professional training postdocs receive (Fig 3B). Finally, these 35 applicants represented 13 different clinical and basic science departments at UMMS. After reviewing the applications, we conducted phone interviews for a small subset to confirm motivation and commitment, which led to the committee inviting 33 postdoctoral fellows to participate with 32 ultimately matriculating into the You^3^ program. Figure 2A illustrates the overall framework for designing and implementing the You^3^ program, which can be adapted as a blueprint by other institutions.

**Figure 3:**
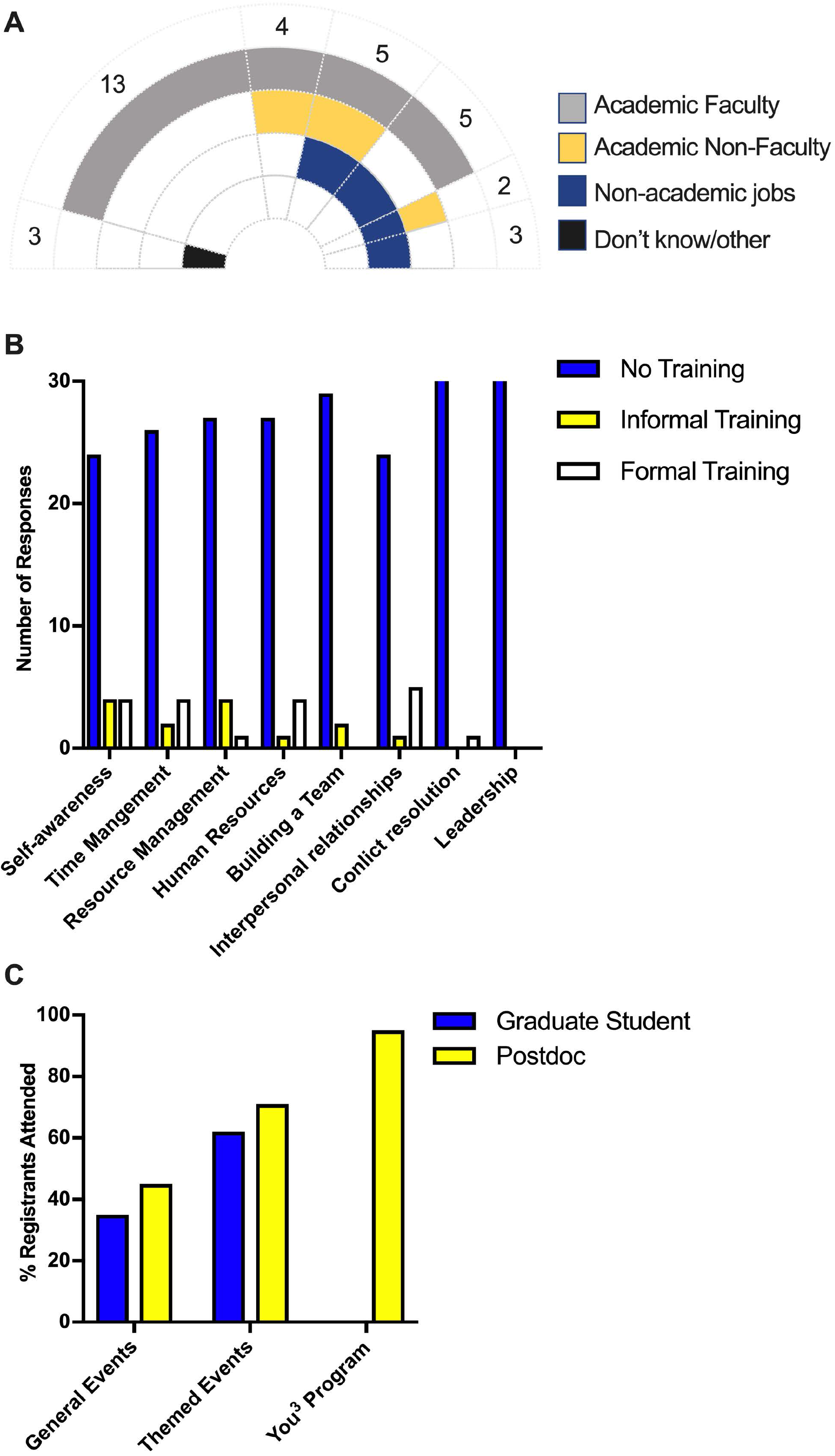
You^3^ participants have diverse career interests but little formal training in professional skills, and they display high commitment to the You^3^ program. (A) Applicants were asked to select one or more career paths they were currently interested in from the broad categories of Academic faculty, Academic non-faculty, Non-academic jobs, and Don’t-Know/Other. 45% of applicants had more than one future career interest. (B) A pre-program anonymous survey asked the You^3^ participants to report any training they had received in Self-awareness, time management, resource management, human resources, building a team, and interpersonal relationships. Their responses were categorized into formal training (led by an instructor with engagement activities, i.e. workshop, class), informal training (purely self-driven, i.e. online webinar, reading, etc.), and no training prior to enrollment. (C) Attendance data was collected from PhD students and postdocs from themed and non-themed events organized by OGPS. The percent registered attendance is compared to the average attendance percentage of the You^3^ participants during the program.

The program consisted of one 90-minute meeting per week for eight consecutive weeks. In addition, there was a welcome event the week prior to the first session and a closing reflection ceremony the week after the last session. Each 90-minute in-class session was led by a subject matter expert external or internal to our institution. The external experts included an academic leadership coach (led four sessions) and a seasoned career consultant (led two sessions). The internal experts included an organizational learning specialist, who led the time management session, and two different unit-level senior administrative leaders, who co-facilitated the session on budgets and finances. The sessions, as such, were a mix of didactic instruction, group activities to enact or respond to scenarios, reflect on, and discuss pertinent topics. Pre-work, outside of class, included short readings and occasional self-assessment material. To further instill structure and weekly learning objectives of the You^3^ program, the program content and calendar reminders were organized on Canvas, a Learning Management System used for courses at our institute. Finally, to facilitate honest conversations and maintain a safe space, there was a community agreement on maintaining confidentiality outside of the class.

Attendance was required for participants in order to receive a certificate of completion. This not only encouraged strong commitment to the program (Fig 3C), but it also enabled sequentially learning. We permitted pre-approved absences and excused attendance under reasonable circumstances. A small number of participants who could not physically be on campus chose the option of watching the program live stream on a video-conferencing platform. You^3^ participants had significantly higher attendance percentages overall compared to other workshops held by our Office of Graduate and Postdoctoral Studies. Specifically, *ad hoc* professional development workshops during the same period had a postdoc attendance of 45% of those who submitted an RSVP, and themed professional development workshops was at 71%. Conversely, the You^3^ program consistently had an attendance record of 95% (Fig 3C).

To measure the impact of the You^3^ program on participants, we decided not to rely on instructor evaluations due to their known biases (13) and inadequacies (14) or on conventional satisfaction surveys that are not meant for learning outcomes. Instead, at the end of our program, we surveyed You^3^ participants as well as non-participating postdocs as a control group to determine self-reported perceptions of a) current knowledge and b) growth during the program time period on each of the eight program topics (refer to Fig S2 for the survey questions). We obtained twice the number of control group respondents as compared to the participants, thus enabling robust comparison while also accounting for inherent biases in self-reporting, such as Dunning-Kruger effect (15). Control group respondents (non-participants) were further stratified between postdocs that have participated in any professional development programming at the University of Michigan and those that did not participate in relevant programming during the same months when the You^3^ program was held.

We used two sets of questions from our survey instrument to measure self-perceptions of a) end-of-program knowledge and b) growth in each module during the duration of the program. This instrument was administered immediately after the program completion to compare knowledge and growth of participants as compared to nonparticipants. The Cronbach’s alpha score was determined for the two primary question blocks stratified by cases and controls (Table 1). These data indicated that our instrument was highly internally reliable.

**Table 1:** Survey Instrument Reliability Analysis. In order to assess survey reliability, Cronbach’s alpha was calculated across relevant sets of questions used to measure current knowledge and growth during program duration. CI= confidence interval.

| **Question Set ↓**<br>**Responder Type→** | **All**<br>**Cronbach's alpha**<br>**Score (95% CI**<br>**range)** | **Participants**<br>**Cronbach's alpha**<br>**Score (95% CI**<br>**range)** | **Control**<br>**Cronbach's alpha**<br>**Score (95% CI**<br>**range)** |
| --- | --- | --- | --- |
| **Growth Question Set** | 0.95 (0.93-0.96) | 0.84 (0.73-0.94) | 0.9 (0.86-0.94) |
| **Current Knowledge Question Set** | 0.83 (0.77-0.88) | 0.83 (0.72-0.93) | 0.78 (0.69-0.87) |

You^3^ participants self-reported greater knowledge at the end of the program on the topics of Hiring and Interviewing, Implicit Bias, Building Your Team, and Budget Management as compared to control groups that had received other professional development programming (Fig 4). Additionally, compared to the no-other-program controls, You^3^ participants reported greater knowledge in all areas with the exception of time management and conflict resolution (Fig 4).

**Figure 4:**
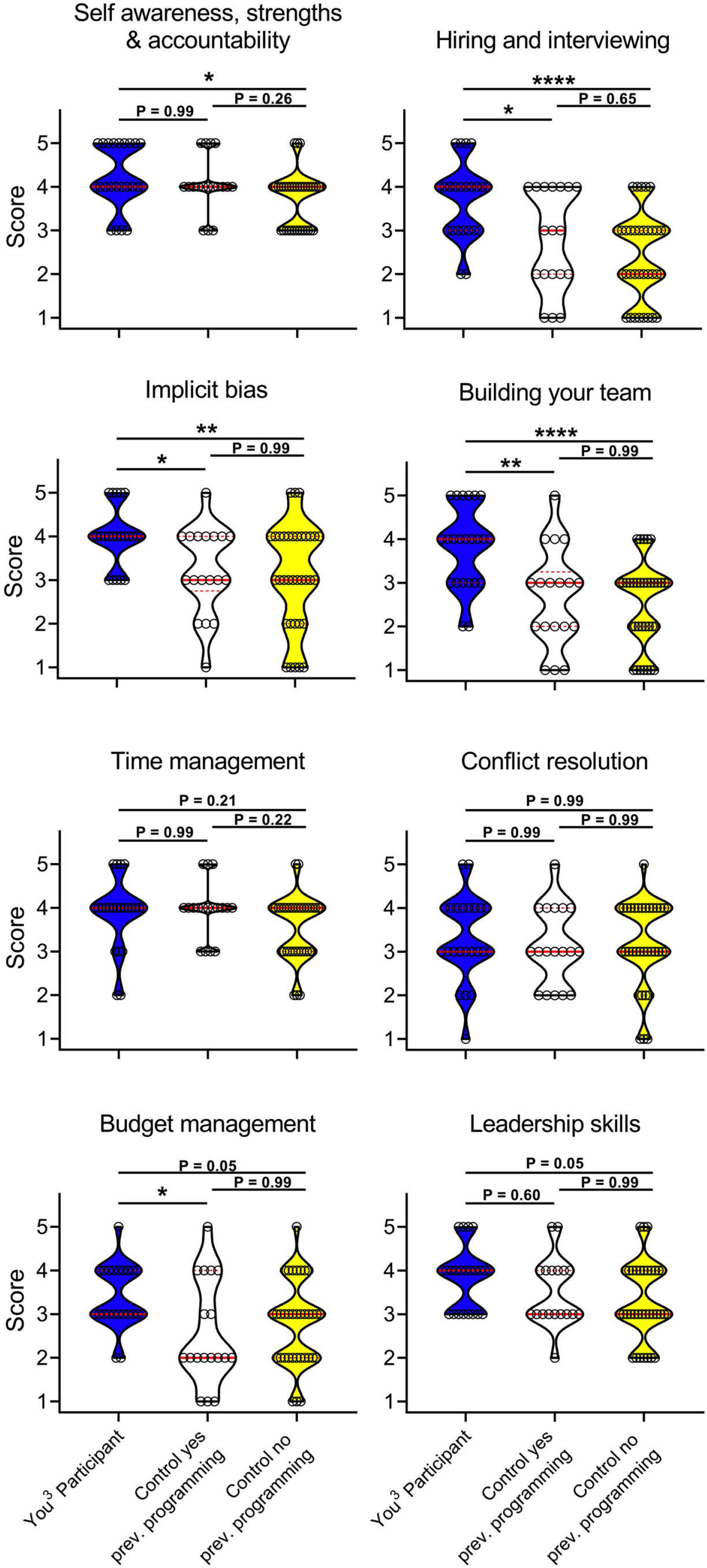
Cross-sectional assessment reveals that You^3^ participants have a higher perception of knowledge than controls across You^3^ program modules. You^3^ participants and controls self-assessed their current knowledge (high = 5, low = 1) for each of the eight You^3^ program modules. Controls were then stratified by previous participation in any career development programming. Current knowledge scores for each module were compared between You^3^ participants (blue, n = 23), controls who had participated in any career development programming (white, n = 18), and controls who had not participated in any career development programming (yellow, n = 35), median (solid red line) and interquartile range (dotted red line) displayed, *P < 0.05, **P < 0.005, ****P < 0.00005, Kruskal-Wallis test followed by Dunn’s posthoc test).

While we cannot quantitatively measure participants’ practical skills in these areas via traditional exams or quizzes, our results indicate the high value You^3^ participants experienced from this program.

To determine if the relationship between comparatively independent, higher self-reported knowledge is associated with participation in the You^3^ program, we also surveyed You^3^ participants and controls on their self-reported growth in each of these skills over the timespan of our program. You^3^ participants reported the highest growth across all skills (Fig 5A). Associative analysis revealed that You^3^ participants had significantly higher odds of reported high growth in 3 of 8 metrics, medium growth in 7 of 8 metrics, and low growth in 1 of 8 metrics (Fig 5B). Therefore, participation in the You^3^ program was highly associated with perceived growth in the topics covered by the You^3^ program. This highlights the value of the You^3^ program as participants feel they have developed greater growth and knowledge in the surveyed topics compared to the control groups.

**Figure 5:**
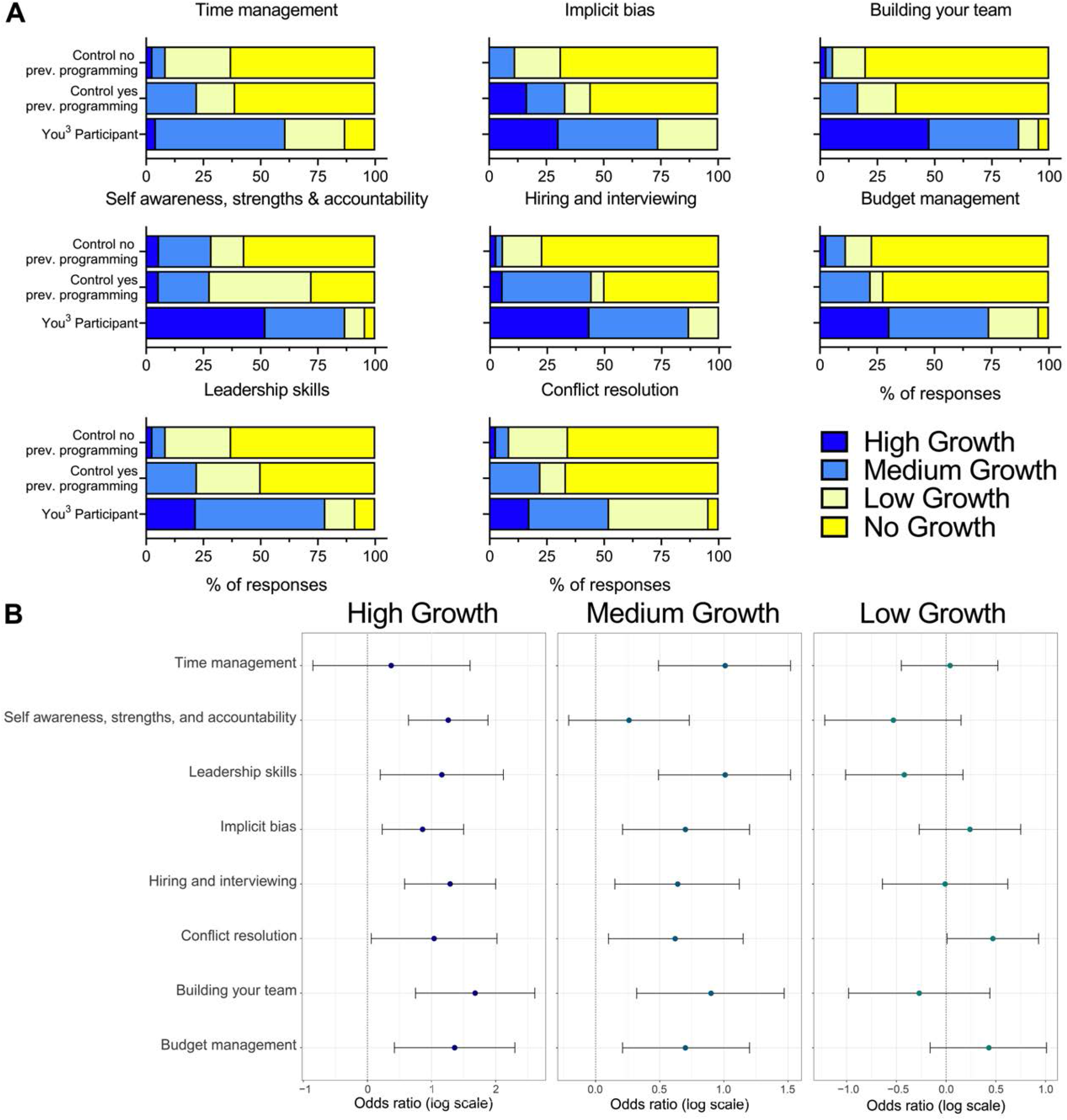
You^3^ participants indicate higher perception of growth across You^3^ program modules compared to controls. You^3^ participants and controls self-assessed their growth (4 = High Growth, 3 = Medium Growth, 2 = Low Growth, 1 = No Growth) over the You^3^ program timeframe for each of the eight You^3^ program modules. Controls were then stratified by previous participation in any career development programming. (A) Growth in each module was compared between You^3^ participants (n = 23), controls who had participated in any career development programming (n = 18), and controls who had not participated in any career development programming (n = 35). (B) Odds ratios for “High Growth,” “Medium Growth,” and “Low Growth” were calculated comparing You^3^ participants (n = 23) to all controls (n = 53) for each of the eight You^3^ program modules (95% confidence intervals displayed).

To see if self-reported knowledge in one topic correlated with knowledge in any of the other seven topics or with years in postdoctoral training, we also performed Pearson’s correlation analyses. We did not find consistent patterns from this analysis when measuring self-reported knowledge correlations for You^3^ participants and previous programming controls. However, we did consistently find statistically significant correlations between all of the eight-knowledge metrics in the no previous programming control group (Fig S3A). These correlations were generally driven by a paucity of self-reported knowledge across our topics, indicating a baseline insufficiency in postdoctoral training related to professional skills. A similar trend was observed when the same correlation analysis was conducted between the self-reported growth metrics, and growth was independent of years in postdoctoral training (Fig S3B). These data indicate that participation in transferable skills development programming is associated with positive growth. Moreover, we posit that the You3 program is more effective at encouraging positive growth than *ad lib* engagement with transferable skills development programming.

**Supplemental Figure 3:**
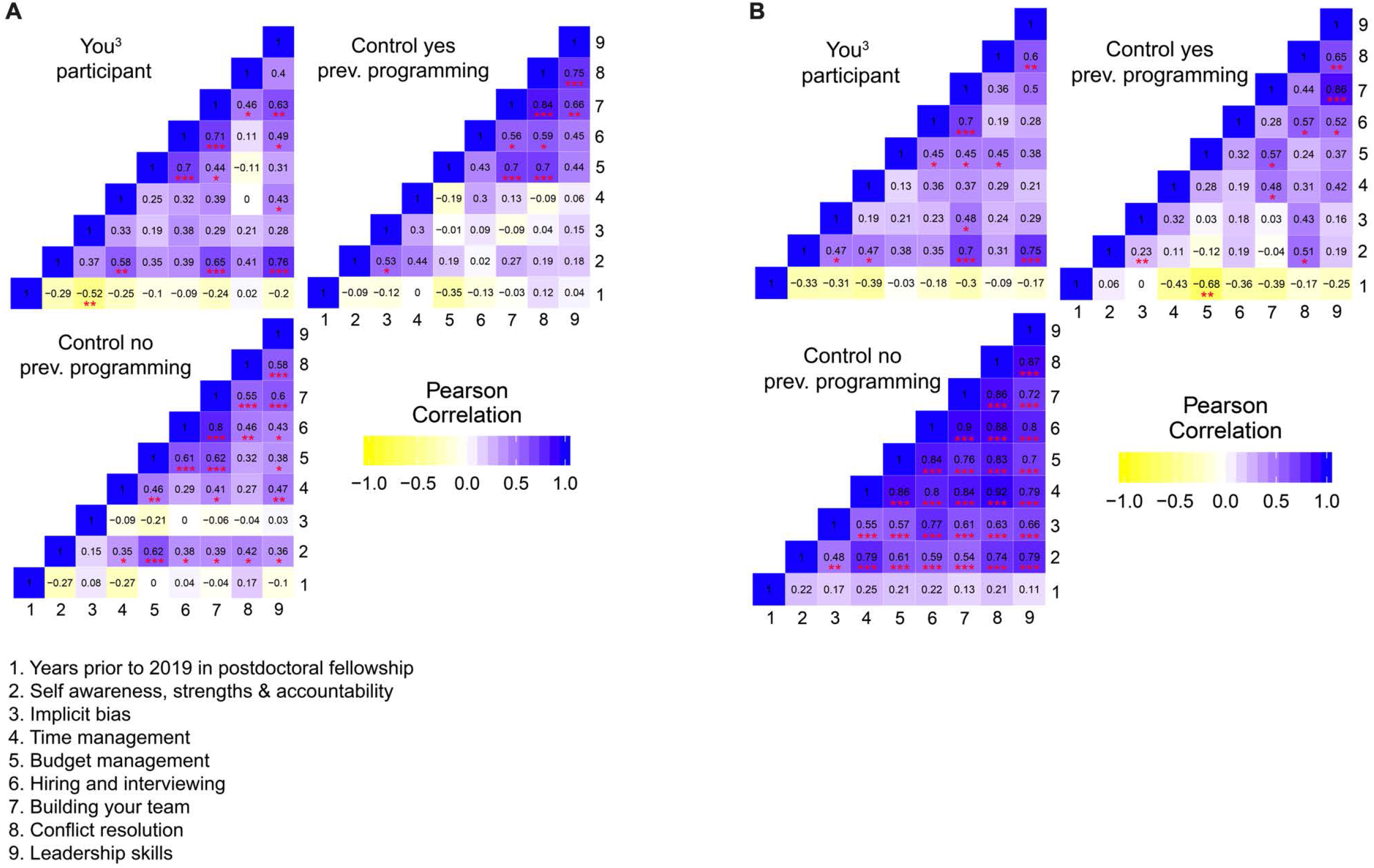
Pearson’s Correlation analysis of self-reported knowledge and growth. (A) Current knowledge scores for each module was correlated to all other modules and the number of years prior to 2019 in their postdoctoral fellowship, and Pearson correlation r and P values were calculated for You^3^ participants, controls who had participated in any career development programming, and controls who had not participated in any career development programming. (B) Growth scores for each module were correlated to all other modules, and Pearson correlation r and P values were calculated for You^3^ Participants, controls who had participated in any career development programming, and controls who had not participated in any career development programming. R values are displayed, and P values are summarized in red (*P < 0.05, **P < 0.005, ****P < 0.0005).

## Discussion

Our study describes a novel program that addresses critical gaps in professional development of postdoctoral fellows. Developed as a collaboration between the Office of Graduate and Postdoctoral Studies (OGPS) and four postdoctoral fellows at the University of Michigan, we structured an array of management and leadership skills that are transferable across careers and assessed program outcomes using rigorous controlled methods. We show that our structured approach resulted in a measurable increase in the participants’ self-reported knowledge and growth in multiple non-technical areas, compared to control non-participants. Interestingly, the increase was independent of time spent as a postdoc, indicating that experience and time alone was not enough to develop skills, and that active programming was necessary.

The unprecedented level of engagement and commitment we catalyzed among the participants as seen by the (a) attendance data and (b) positive outcomes determined by our controlled study could both be due to two key aspects of the program. The first is that the program was fully structured and sequential, which is uncommon for professional skill development activities for postdocs. The opportunities routinely available for skill building for trainees tend to be independent workshops that do not have the pedagogical structure and continuity necessary for comprehensive learning (9) This lack of structure is a problem especially for postdocs, as they have a short time frame to be productive and to transition from being a trainee to an independent professional (1,8,17). The structure we provided, based on data generated from surveys of the whole academic community as well as future employer feedback and published literature, likely spurred engagement of the participants. The second key innovation was the extent of input from and involvement of four current postdocs, the target population, in designing the program. This is a notable difference between You^3^ and most previous programs. An additional benefit of involving trainees during program development was that the postdocs who helped develop the program also grew as leaders and strategic thinkers. These benefits are consistent with previous reports where engaging college students early has been valuable in developing teaching methods and planning curricula (18).

The novel method that we developed for assessing outcomes, modeled on the scientific method, is a key advance. Traditionally, educational and professional development programs rely on satisfaction surveys and/or instructor ratings, both of which are beset with issues such as bias and lack of depth (19, 20). Attempts to assess outcomes in more quantitative ways have been challenging, as these programs typically do not (and should not) use grades as a measurement of learning. The controlled assessments, comparing participants’ perceived knowledge and growth to non-participants’ in the same time frame, was developed based on a method used in medical settings such as clinical trials. Because self-assessment and awareness form a significant part of the assessments, it is possible that the assessments were influenced by participants’ awareness of what they did not know earlier.

The outcomes and the success of the inaugural offering as described in this manuscript has informed us to continue this program, while making it stronger for future offerings. But due to the sudden changes to in-person gatherings and due to significant budget cuts, our emphasis for the next cohort was to optimize the program for virtual and hybrid formats during the COVID-19 pandemic. Albeit virtual, based on feedback from the 2019 cohort, we implemented a greater number of activities and assignments to be completed during and outside of the session. While we are still collecting feedback from these changes, our goal was to increase experiential learning and build community among postdocs who often suffer from isolation and loneliness (21,22). Our 2020-21 experience, along with another recent study our office conducted, unearthed several advantages of virtual professional development programming, prompting us to likely keep a virtual component in some form (23).

An aspect of the program that we plan to develop and build on is long-term tracking of the postdocs who have participated in the program. We anticipate gathering data on not only their professional paths but also intend to set up mechanisms to best understand how this program set the foundation for lifelong learning in critical areas such as emotional intelligence (24)(25) and related interpersonal skills. This longitudinal data, along with information on how other institutions are implementing similar programs, will strengthen our assessment. Using this iterative approach, we aim to optimize the program to its most effective format.

The You^3^ program is a novel, scalable, and rigorous framework that can be adapted by any institution to expand the foundation of postdoctoral training. Our structured program and scientific assessment of outcomes helps minimize the variabilities and inequities in experiences and consequences prevalent in the current apprenticeship-based training models. For example, postdocs don’t have equitable time or guidance to access professional skill building and career development, despite recommendations from national bodies like the National Postdoc Association and relevant studies (16)(2). A subset of postdocs who are independently funded, such as by the NIH’s K or F award systems, require professional development plans to be submitted in the proposal and followed through. On the contrary, postdocs who are not independently funded don’t have any mechanism to consider professional development. Structured programs such as You^3^, developed via postdoc-centric guiding principles, mitigate such inequities by providing access to professional development in a structured manner to all postdocs regardless of funding status. Our program is an important benefit for international postdocs, who often don’t get the opportunity to craft plans via funding proposals, as they are often ineligible for independent funding.

Finally, based on our experience, we believe You^3^ can be adapted easily to institutional needs, including hybrid training models combining in-person and virtual formats. We anticipate that newly funded initiatives from the NIH, such as The Postdoc Academy (<u>postdocacademy.org</u>), will provide complementary and supplementary resources to make broad implementation feasible within the budgets of most institutions.

## Acknowledgments

We are grateful to Dr. Manoj Puthenveedu & Dr. Michele Swanson for critical reading and feedback on the manuscript. We thank instructors and content contributors to the program. We acknowledge, as part of a larger consulting project for our office, the efforts of the miLEAD team. Finally, we are thankful to Associate Dean Dr. Mary O’Riordan, Dr. Michele Swanson and members of the Office of Graduate and Postdoctoral Studies for being supportive of our work. Dr. Shoba Subramanian is partly funded by an NSF-IGE grant (#1954967).

---

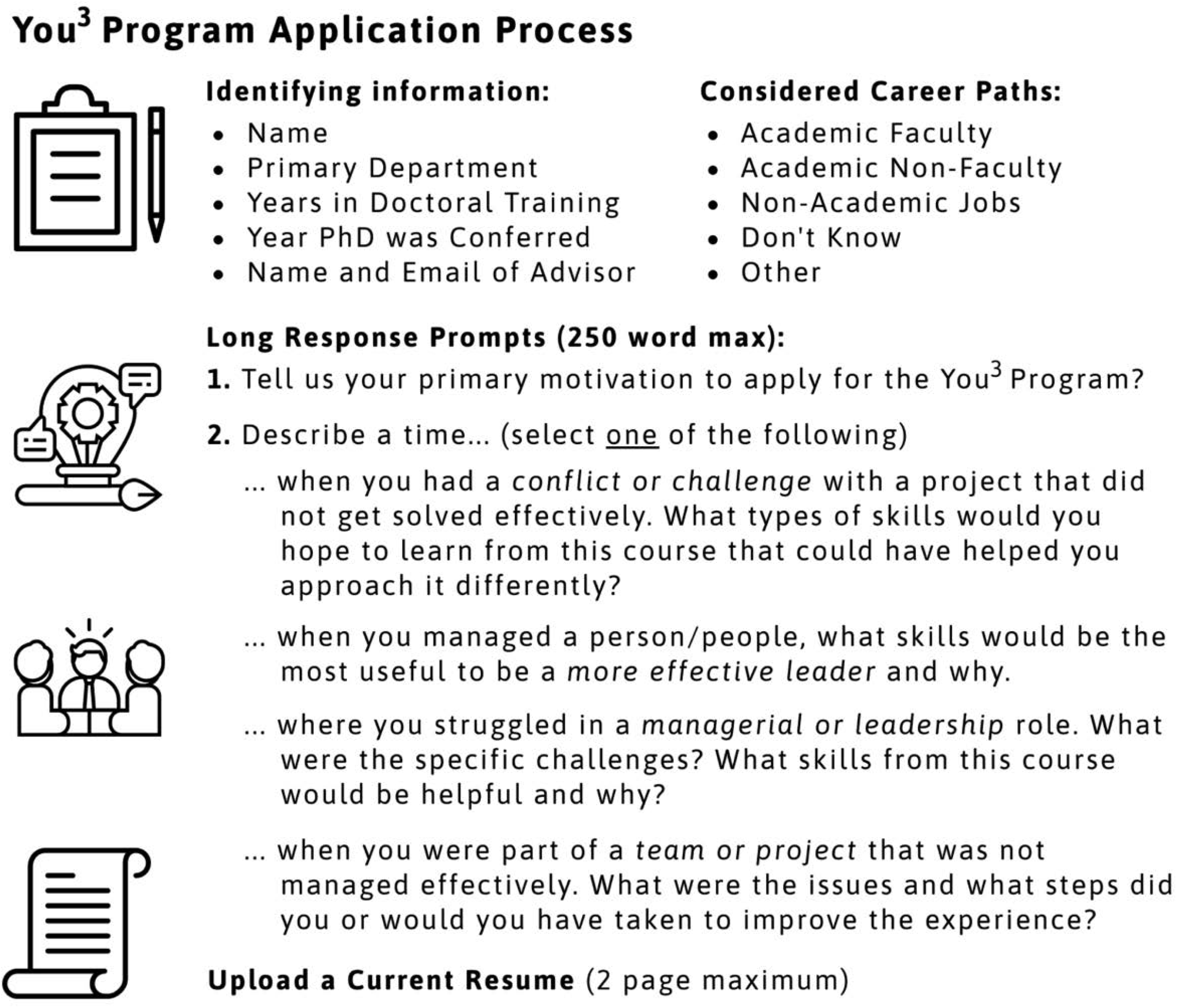

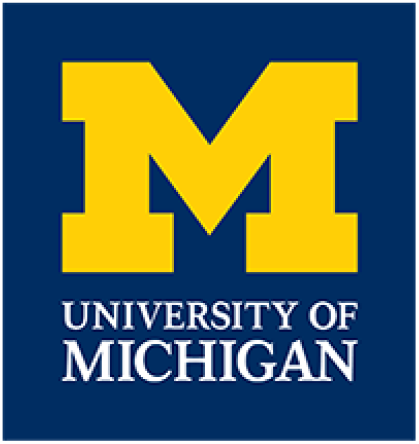

## Block 3

We’re inviting you to participate in a research study to enhance current and future professional development programs for postdoctoral fellows at the University of Michigan. Participation is completely voluntary and response to any survey questions indicates your consent. Data collection is anonymous, however; once the survey is initiated (answering a question), your data cannot be removed. There are no negative consequences to participation, whatever you decide. If you have any questions or concerns, please contact Shoba Subramanian at shobas@umich.eduor734-615-6511.

## Block 1

Select the year when you started as a postdoc at the University of Michigan

∘ 2014
∘ 2015
∘ 2016
∘ 2017
∘ 2018
∘ 2019

Measuring Participants’ Knowledge and Growth for Each of the You3 Topics

Measuring Participants' Knowledge and Growth for Each of the You3 Topics
|  | Rate on a scale of 1-5 your<br>**\*\*current\*\* end-of-program**<br>knowledge in each of the You3 topic<br>modules (1: very low; 3: medium; 5:<br>very high) |  |  |  |  | Document any **increase in knowledge**<br>**as a result the You3 program**<br>participation. (Calibrate your baseline to<br>knowledge before program started) |  |  |  |
| --- | --- | --- | --- | --- | --- | --- | --- | --- | --- |
|  | 1 | 2 | 3 | 4 | 5 | No<br>Growth | Low<br>Growth | Medium<br>Growth | High<br>Growth |
| Self awareness, strengths & accountability | ☐ | ☐ | ☐ | ☐ | ☐ | ☐ | ☐ | ☐ | ☐ |
| Implicit bias | ☐ | ☐ | ☐ | ☐ | ☐ | ☐ | ☐ | ☐ | ☐ |
| Time management | ☐ | ☐ | ☐ | ☐ | ☐ | ☐ | ☐ | ☐ | ☐ |
| Budget management | ☐ | ☐ | ☐ | ☐ | ☐ | ☐ | ☐ | ☐ | ☐ |
| Hiring, Interviewing | ☐ | ☐ | ☐ | ☐ | ☐ | ☐ | ☐ | ☐ | ☐ |
| Building your team | ☐ | ☐ | ☐ | ☐ | ☐ | ☐ | ☐ | ☐ | ☐ |
| Conflict resolution | ☐ | ☐ | ☐ | ☐ | ☐ | ☐ | ☐ | ☐ | ☐ |
|  | 1 | 2 | 3 | 4 | 5 | No<br>Growth | Low<br>Growth | Medium<br>Growth | High<br>Growth |
| Leadership<br>skills | ☐ | ☐ | ☐ | ☐ | ☐ | ☐ | ☐ | ☐ | ☐ |

Measuring applicability of knowledge gained from You3 program participation. The application can be for a current or potential area of concern and can be in current postdoc position or in a career after your postdoc training.

|  | I plan to use the below skills<br>& knowledge learned in You3<br>in a future professional<br>space (current position or<br>new career). |  | Provide 1-2 examples of where you will/could use<br>knowledge from this module. If you answered "no",<br>then say n/a. Example: Lab mentoring, ongoing<br>collaboration, potential conflict, planning job<br>applications, etc. |
| --- | --- | --- | --- |
|  | Yes | No | Type 1-2 examples here |
| Self<br>awareness,<br>strengths &<br>accountability | ☐ | ☐ |  |
| Implicit bias | ☐ | ☐ |  |
| Time<br>management | ☐ | ☐ |  |
| Budget<br>management | ☐ | ☐ |  |

|  | I plan to use the below skills & knowledge learned in You3 in a future professional space (current position or new career). |  | Provide 1-2 examples of where you will/could use knowledge from this module. If you answered "no", then say n/a. Example: Lab mentoring, ongoing collaboration, potential conflict, planning job applications, etc. |
| --- | --- | --- | --- |
|  | Yes | No | Type 1-2 examples here |
| Hiring, Interviewing | ☐ | ☐ |  |
| Building your team | ☐ | ☐ |  |
| Conflict resolution | ☐ | ☐ |  |
| Leadership skills | ☐ | ☐ |  |

List any other workshop** you participated in between Sept and Nov 2019. Note: A workshop (in this context) is defined as a 1-hour or longer in-person program on a professional development topic led by an expert.

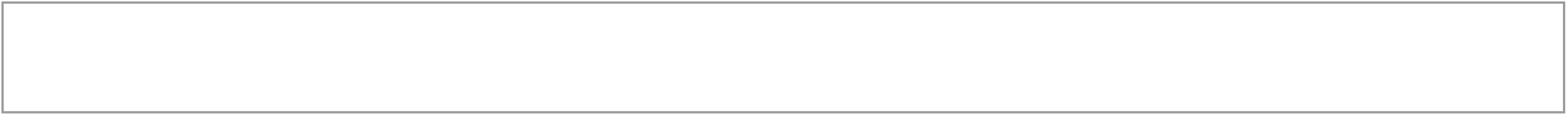

## You3 Program Evaluation

Overall, how satisfied or dissatisfied were you with the You3 Program?

∘ Extremely satisfied
∘ Moderately satisfied
∘ Slightly satisfied
∘ Neither satisfied nor dissatisfied
∘ Slightly dissatisfied
∘ Moderately dissatisfied
∘ Extremely dissatisfied

How likely are you to recommend this course to a friend or classmate?

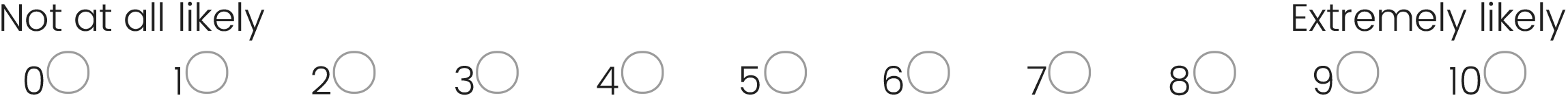

How much did you learn from this course?

∘ A great deal
∘ A lot
∘ A moderate amount
∘ A little
∘ Nothing at all

How reasonable or unreasonable was the time allocated for this course?

∘ Extremely reasonable
∘ Moderately reasonable
∘ Slightly reasonable
∘ Neither reasonable nor unreasonable
∘ Slightly unreasonable
∘ Moderately unreasonable
∘ Extremely unreasonable

How knowledgeable was the instructor of the material presented in this course?

∘ Extremely knowledgeable
∘ Very knowledgeable
∘ Moderately knowledgeable
∘ Slightly knowledgeable
∘ Not knowledgeable at all

How well did this course meet your expectations?

∘ Extremely well
∘ Very well
∘ Moderately well
∘ Slightly well
∘ Not well at all

What did you like most about this program?

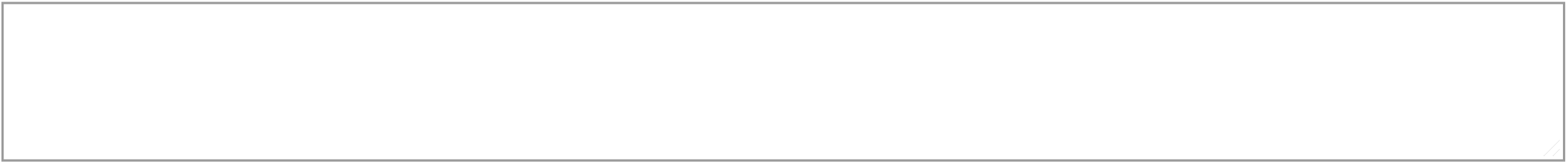

What did you like least about this program?

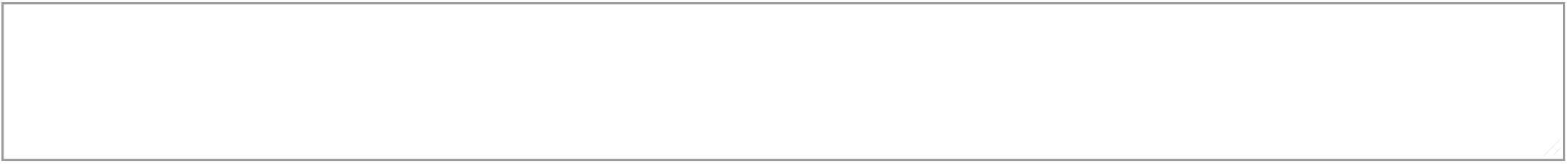

How could this program be improved?

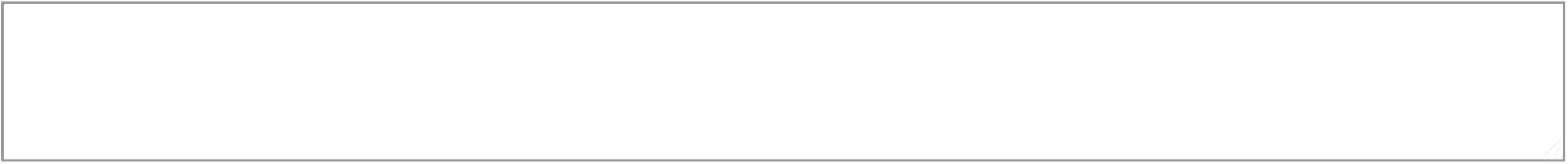

## Block 3

Thank you for participating in this inaugural program and for filling out the survey!

<u>Powered by Qualtrics</u>

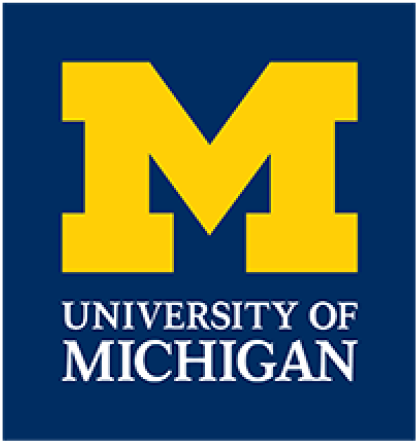

## Purpose of Survey

We’re inviting you to participate in a research study to enhance current and future professional development programs for postdoctoral fellows at the University of Michigan. Participation is completely voluntary and response to any survey questions indicates your consent. Data collection is anonymous, however; once the survey is initiated(answering a question), your data cannot be removed. There are no negative consequences to participation, whatever you decide. If you have any questions or concerns, please contact Shoba Subramanian at 734-615-6511.

## Participation in You3

Have you heard of the <u>You3 Leadership & Management Program</u> for Postdocs at the University of Michigan Medical School?

∘ Yes
∘ Maybe
∘ No

Did you participate in the OGPS You3 Program - Sept-Nov 2019?

∘ Yes
∘ No

Select the year when you started as a postdoc at the University of Michigan

∘ 2014
∘ 2015
∘ 2016
∘ 2016
∘ 2017
∘ 2018
∘ 2019

## Measure You3 Topic Knowledge & Growth

Measuring Knowledge and Growth for Leadership & Management (as they apply in a professional work place).

|  | Rate on a scale of 1-5 your **\*\*current\*\* knowledge in each of the topics** (1: very low; 3: medium; 5: very high) |  |  |  |  | Document any **increase in knowledge (within the last 3 months)** based on participation in any recent workshops |  |  |  |
| --- | --- | --- | --- | --- | --- | --- | --- | --- | --- |
|  | 1 | 2 | 3 | 4 | 5 | No Growth | Low Growth | Medium Growth | High Growth |
| Self awareness, strengths & accountability | ☐ | ☐ | ☐ | ☐ | ☐ | ☐ | ☐ | ☐ | ☐ |
| Implicit bias | ☐ | ☐ | ☐ | ☐ | ☐ | ☐ | ☐ | ☐ | ☐ |
| Time management | ☐ | ☐ | ☐ | ☐ | ☐ | ☐ | ☐ | ☐ | ☐ |
| Budget management | ☐ | ☐ | ☐ | ☐ | ☐ | ☐ | ☐ | ☐ | ☐ |
| Hiring, Interviewing | ☐ | ☐ | ☐ | ☐ | ☐ | ☐ | ☐ | ☐ | ☐ |
| Building your team | ☐ | ☐ | ☐ | ☐ | ☐ | ☐ | ☐ | ☐ | ☐ |
| Conflict resolution | ☐ | ☐ | ☐ | ☐ | ☐ | ☐ | ☐ | ☐ | ☐ |
| Leadership skills | ☐ | ☐ | ☐ | ☐ | ☐ | ☐ | ☐ | ☐ | ☐ |

List any workshops that you participated in between Sept and Nov 2019. Note: A workshop (in this context) is defined as a 1-hour or longer in-person program on a professional development topic led by an expert.

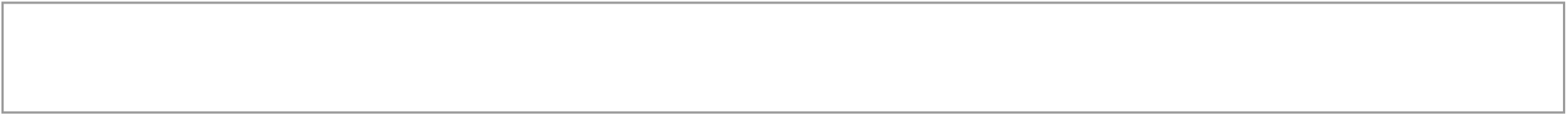

## You3 Program Evaluation

If offered next year, will you apply to participate in the You3 Postdoc Leadership and Management Program?

<u>You3 Program Link</u>

∘ Extremely likely
∘ Somewhat likely
∘ Neither likely nor unlikely
∘ Somewhat unlikely
∘ Extremely unlikely

Optional: If you are unable to participate in You3, what are the barriers to this?

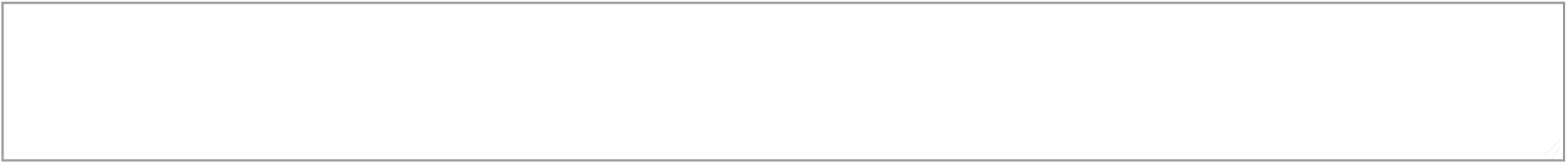

Optional: What other professional development workshops will you attend, if offered? Please list topics.

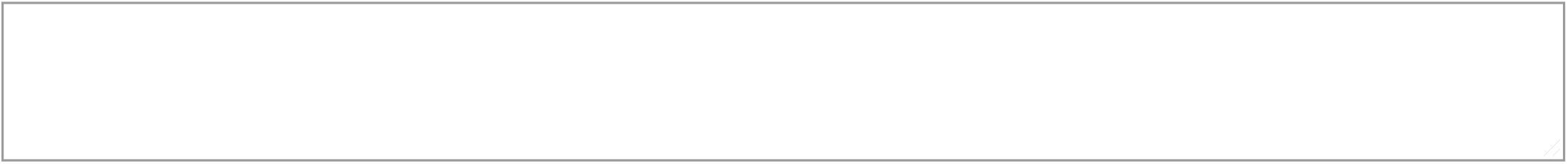

## Block 4

Thank you for clicking on this survey, since you participated in You3, please respond to the survey posted via the Canvas site tailored for program participants

Thank you for filling out this survey!

